# Single-cell Analysis of Attenuation-Driven Transcription Reveals New Principles of Bacterial Gene Regulation

**DOI:** 10.1101/2025.10.06.680652

**Authors:** S Moradian, N Ali, R Jagadeesan, V Behrends, AS Ribeiro, C Engl

**Affiliations:** School of Biological and Behavioural Sciences, Queen Mary University of London, UK; Laboratory of Biosystems Dynamics, Faculty of Medicine and Health Technology, Tampere University, Finland; School of Medicine and Biosciences, University of West London. UK; Department of Biochemistry, University of Cambridge, UK

## Abstract

Transcription attenuation fine-tunes biosynthetic gene expression in bacteria via premature termination upon metabolic signals. In transcription initiation-controlled bacterial systems, promoter architecture and transcription factor binding sets the size of transcriptional bursts at σ^70^ promoters, while distal enhancer elements and associated transcriptional activators modulate burst frequency at σ^54^ promoters. Using the tryptophan biosynthesis operon as a model, we show that transcription attenuation, acting post-initiation and alongside transcriptional repression, simultaneously modulates both burst size and frequency from a σ^70^ promoter. This challenges the view that frequency modulation requires distal enhancer input and reveals that post-initiation mechanisms can shape divergent transcriptional bursting. We also uncover that bacteria use cross-feeding as a previously unrecognised strategy for controlling cell-to-cell variation in gene expression, with implications for metabolic coordination among cells. These findings redefine transcription dynamics within cell populations and suggest new principles by which bacteria regulate gene expression to adapt to environmental change.

## Introduction

In both prokaryotes and eukaryotes, transcriptional output often occurs in bursts, short episodes of mRNA synthesis interrupted by periods of transcriptional silence (1–8). This transcriptional bursting gives rise to significant cell-to-cell variability in mRNA levels, even among genetically identical individuals in homogeneous environments (5–10). Such noise in gene expression facilitates phenotypic heterogeneity, enabling bacterial subpopulations to survive unfavourable conditions such as antibiotic exposure or nutritional depletion (11,12).

Recent studies have made considerable progress in linking transcriptional bursting dynamics to promoter architecture and transcription initiation mechanisms (2,10,13–21). However, post-initiation regulatory mechanisms, such as the premature termination of transcription, remain underexplored in this context, particularly regarding how they influence bursting kinetics at the single-cell level. Addressing this gap is critical to better understand the relationship between gene regulation mechanisms and cellular adaptations, e.g. in response to treatment. In these conditions, many biosynthetic operons including those responsible for amino acid and nucleotide production, are subject to premature termination of transcription (22,23).

A classic model for premature termination of transcription is the tryptophan (*trp*) operon, which regulates tryptophan biosynthesis (24–29). Despite the conservation of the operon across species, *E. coli* and *B. subtilis* have evolved distinct regulatory mechanisms (30) (Fig. 1). In *E. coli*, the *trp* operon is regulated by both repression at the level of transcription initiation via the TrpR repressor and attenuation, acting at the post-initiation stage, through the TrpL leader peptide-mediated formation of a transcriptional terminator (29,30). By contrast, *B. subtilis* primarily relies on post-initiation attenuation mediated by TRAP (*trp* RNA-binding attenuation protein), without a transcriptional regulator acting at initiation (24,31–34). These differing regulatory strategies provide an ideal comparative model to investigate how single-layer (post-initiation only, PI) versus dual-layer (initiation and post-initiation, I+PI) control influences transcriptional noise and bursting behaviour.

**Fig. 1.**
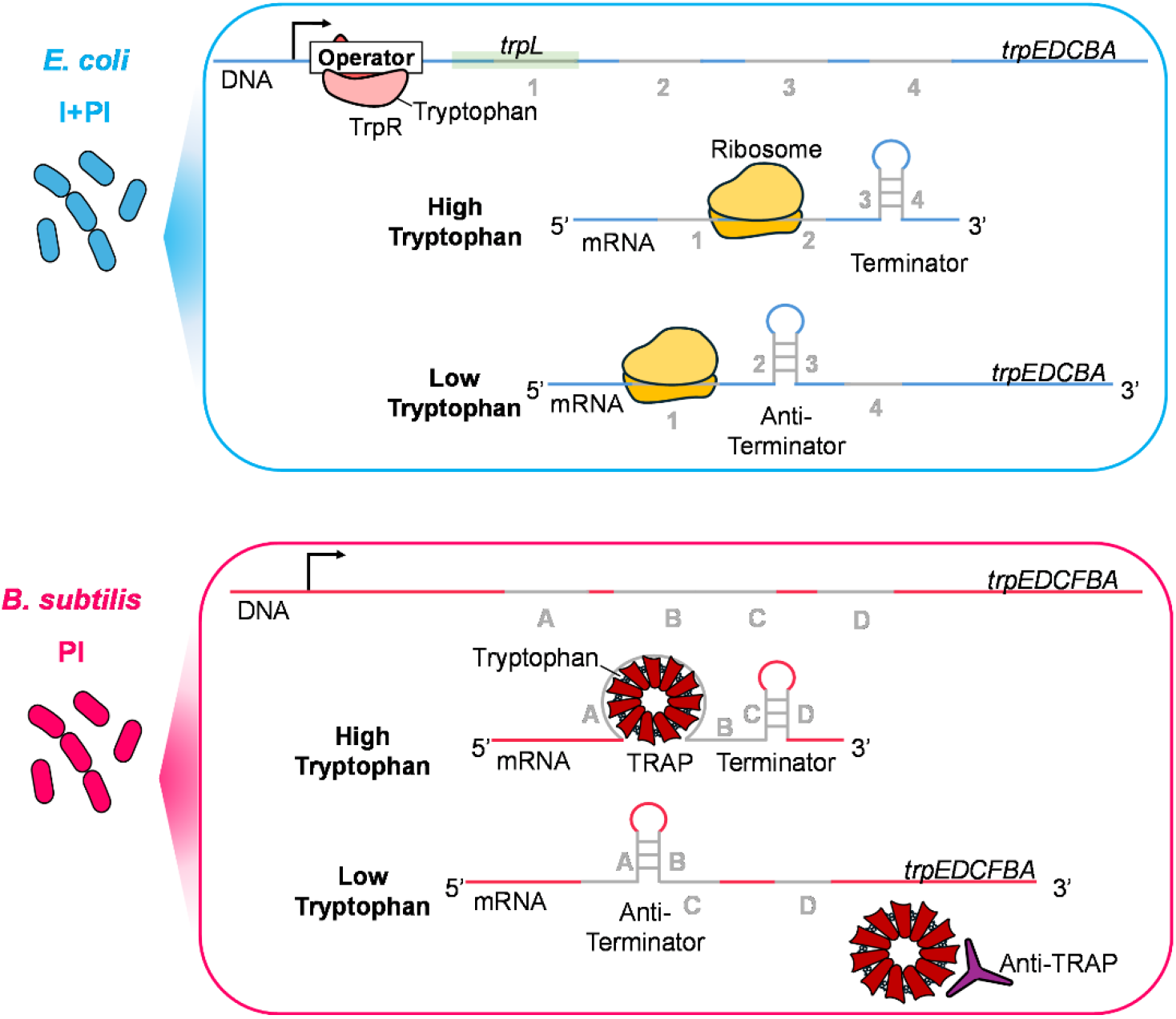
Dual-verse single-layer transcription regulation strategies of the *trp* operon. Schematic of the dual-layer (I+PI) regulation in *E. coli* and the post-initiation–only (PI) regulation in *B. subtilis*. In *E. coli*, tryptophan-activated TrpR inhibits transcription initiation by RNA polymerase (RNAP) through binding to the operator region upstream of the structural genes. At the post-initiation level, high levels of charged tRNA^Trp^ promote rapid ribosome translation of *trpL* region 1, which favours formation of the terminator (3:4) hairpin, leading to transcriptional attenuation. Under tryptophan limitation, TrpR remains inactive, allowing RNAP to initiate transcription, and ribosome stalling at region 1 due to reduced charged tRNA^Trp^ availability enables formation of the antiterminator hairpin (2:3), resulting in transcriptional readthrough of the operon. In *B. subtilis*, RNAP initiates transcription constitutively, and regulation occurs solely at the post-initiation stage. Tryptophan-activated TRAP binds to the leader mRNA to promote terminator hairpin (C:D) formation which prematurely terminates transcription. Under tryptophan limitation, anti-TRAP binds with tryptophan activated TRAP to inhibit its RNA-binding ability, preventing premature termination, thereby allowing the antiterminator (A:B) to form and transcription to proceed into the downstream structural genes of the operon.

In both organisms, the *trp* operons are transcribed from housekeeping sigma factors, σ^70^ in *E. coli* and σ^A^ (a member of the σ^70^ family) in *B. subtilis*. In bacteria, σ^70^-dependent promoters drive burst-size modulated transcription when regulated at initiation, producing mRNA in episodic bursts (2,13). Transcriptional noise in such promoters, however, often decreases at higher expression levels (9,13,14). To date, the effect of pre-mature termination by transcription attenuation on noise and bursting in these promoters remains poorly understood and is the focus of this study (2,13).

## Results

### Single-layer transcriptional regulation by attenuation allows persistence of high-expressing subpopulations

To compare the effects of post-initiation-only regulation versus a dual-layered mechanism involving both transcription initiation and post-initiation control, on transcriptional activity, we used wildtype (WT) strains of *B. subtilis* and *E. coli*, referred to throughout this article as PI (post-initiation only) and I+PI (initiation plus post-initiation), respectively (35,36). Transcriptional heterogeneity and burstiness were measured across populations exposed to varying concentrations of tryptophan: 0, 5, 50, and 100 µM, denoted as 0, 1X, 10X, and 20X, respectively. We used RNA FISH (37) to quantify *trpE* mRNA expression at single-cell and single-molecule resolution.

To assess how regulatory strategies shape transcript output at the single-cell level, we quantified *trpE* mRNA copy-number distributions (Fig. 2A). Under tryptophan starvation, both modes showed heterogeneous expression, but the maximum output differed: I+PI cells produced up to 27 transcripts, while PI-only reached 49. This suggests that the additional layer of repression at transcription initiation in the I+PI system not only reduces the proportion of transcriptionally active cells but also constrains their transcriptional output capacity. With added tryptophan, I+PI cells maintained only 1–2 transcripts, eliminating high expressors from the population, whereas PI-only retained a subpopulation of cells with more than 5 transcripts even at 20X tryptophan. These pre-adapted cells may provide a survival advantage at the population level during abrupt depletion of environmental tryptophan.

**Fig. 2.**
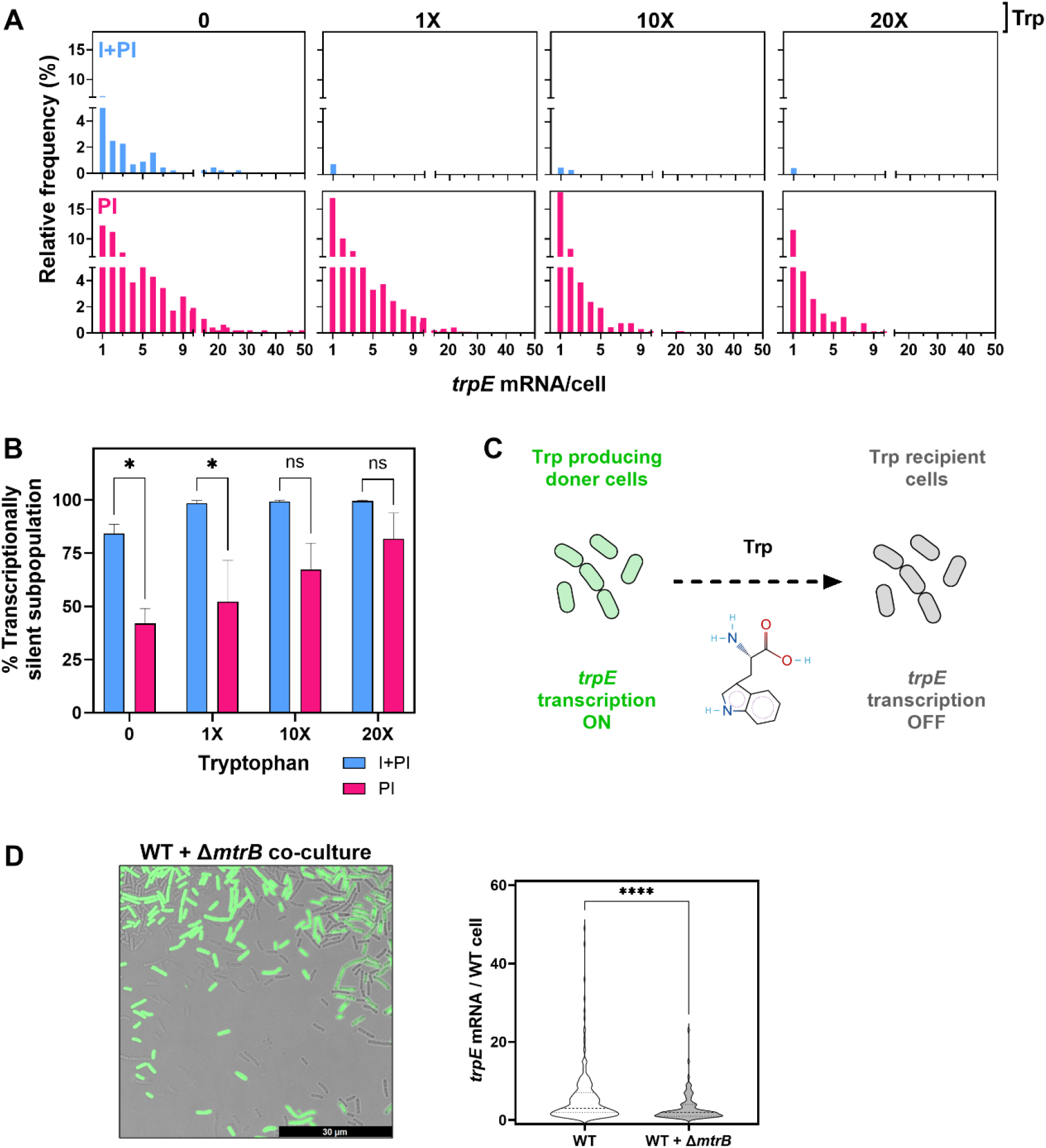
Impact of single- and dual-layer transcriptional regulation on *trpE* transcriptional heterogeneity. I+PI (*E. coli*) and PI (*B. subtilis*) cells were cultured with 0, 1X, 10X, or 20X tryptophan (0, 5, 50, and 100 µM), and single-cell *trpE* mRNA counts were obtained using smRNA FISH. **(A)** Frequency distributions of *trpE* transcripts in transcriptionally active cells (≥1 mRNA). **(B)** Percentage of transcriptionally silent cells (0 mRNA). Asterisks indicate statistical significance assessed by two-way ANOVA followed by Tukey’s multiple comparisons test (GraphPad Software Inc.). **(C)** Model proposing that tryptophan cross-feeding results in inconsistencies in expression of biosynthetic genes: active cells synthesise tryptophan and provide it to recipient cells, which then downregulate *trpE* transcription, creating heterogeneity. **(D)** Left panel: fluorescence images of WT and GFP-labelled Δ*mtrB* cocultures. *trpE* mRNA molecules were quantified in WT cells grown either alone or with the high *trpE*-expressing Δ*mtrB* mutant. Right panel: violin plots showing single-cell *trpE* distributions in monoculture versus coculture, in cells with ≥1 *trpE* transcript. The plots demonstrate variability and overall distribution of *trpE* expression. A Mann–Whitney test was performed using GraphPad Prism to assess statistical significance (p < 0.0001).

Cells were characterised as transcriptionally silent if they did not contain any *trpE* mRNA molecules. Interestingly, even in the absence of any exogenously supplemented tryptophan, only a subset of cells exhibited *trpE* expression (Fig. 2B) This persistence of a transcriptionally silent subpopulation under tryptophan-starved conditions was observed in both regulatory modes, but the proportion of inactive cells was double in the dual-layered regulatory system (I+PI), where 84% of cells were inactive, compared to 42% in the single-layer post-initiation (PI) model.

Statistical analysis confirmed that the differences in the percentage of transcriptionally inactive cells between the I+PI and PI regulation modes were significant at both 0X and 1X tryptophan conditions (p = 0.026 and p = 0.019 respectively). As extracellular tryptophan concentrations increased, both modes exhibited an increase in the proportion of transcriptionally inactive subpopulation. However, the magnitude and sensitivity of this response is more pronounced in the I+PI mode, indicating that dual-layered regulation provides a greater capacity to shift transcriptional activity across the population in response to metabolic cues. In contrast, the PI system showed a more gradual, dose-dependent increase in transcriptional inactivation.

### High trpE-expressing cells suppress trpE transcription in recipient cells, promoting heterogeneity

The persistence of *trpE* non-transcribing cells during tryptophan starvation, shown in Fig. 2B, is unexpected, given the amino acid’s essential role in growth. One explanation is uptake of tryptophan produced by neighboring *trpE*-active cells. We therefore asked whether high *trpE* expressors (here labelled with GFP to discriminate from WT cells) could suppress transcription in recipient WT cells, allowing a silent subpopulation to persist even without external tryptophan (Fig. 2C).

First, we asked whether tryptophan could be released by *trpE*-expressing producer cells and taken up by auxotrophs lacking *trpE* (Fig. S1). Extracellular amino acids were measured in *E. coli* and *B. subtilis* WT cultures grown in minimal media, revealing the presence of tryptophan in supernatants of both species (Table S1). This is unlikely to result from cell lysis: intracellular tryptophan in *E. coli* is <1% of aspartate, yet abundant intracellular amino acids such as aspartate, lysine, and arginine were absent from the supernatant, indicating selective metabolite release. Other costly amino acids, including phenylalanine and tyrosine, were also detected, suggesting cross-feeding of metabolically expensive compounds. Growth of Δ*trpE* auxotrophs on WT spent medium, but not fresh minimal medium, confirmed that tryptophan was released and available for uptake (Fig. S2). Together, these data demonstrate that *trpE* producer cells can release tryptophan into the extracellular environment and subsequently taken up by non *trpE* producer cells, suggesting the possibility of cross-feeding.

Strikingly, our data further suggests that cross-feeding can act as a regulatory mechanism of gene expression enabling transcriptional crosstalk between cells. Using *B. subtilis* co-culture as a model, we found that the presence of high *trpE*-expressing Δ*mtrB* cells (lacking transcription attenuation protein TRAP) suppresses *trpE* transcription at the native chromosomal locus of WT cells (Fig. 2D). Single-cell analysis revealed a 46% decrease in mean *trpE* mRNAs per WT cell in co-culture with Δ*mtrB*. The fraction of transcriptionally silent cells increased by 12%, while high (>5 mRNAs) and ultra-high (>30 mRNAs) expressing WT cells were reduced or eliminated (Fig. S3). This suppression in WT cells likely reflects enhanced tryptophan supply by Δ*mtrB* cells, which triggers TRAP-mediated attenuation of *trpE* transcription in WT cells.

### Dual-layer transcriptional regulation drives abrupt shifts in noise and burst

To assess how post-initiation mechanisms combined with repression at initiation shape transcriptional noise, we quantified *trpE* noise in the I+PI (dual-layer) and PI (single-layer) modes.

Tryptophan supplementation increased noise in both modes, but I+PI consistently showed higher noise across all concentrations, including 0X tryptophan, indicating that dual regulation inherently promotes greater noise in gene expression regardless of environmental tryptophan availability (Fig. 3A). In PI, noise increased gradually with tryptophan concentrations, whereas in I+PI noise remained relatively high across the same range. Linear regression revealed moderate logarithmic relationships between expression levels and noise in both modes (R^2^ = 0.66). Spearman’s analysis showed a strong negative correlation between mean *trpE* expression and noise (*ρ* = –1.0, p = 0.08). In both systems, noise declined as expression increased, but the pattern differed: PI showed a gradual reduction, whereas I+PI displayed a sharper, more abrupt drop.

**Fig. 3.**
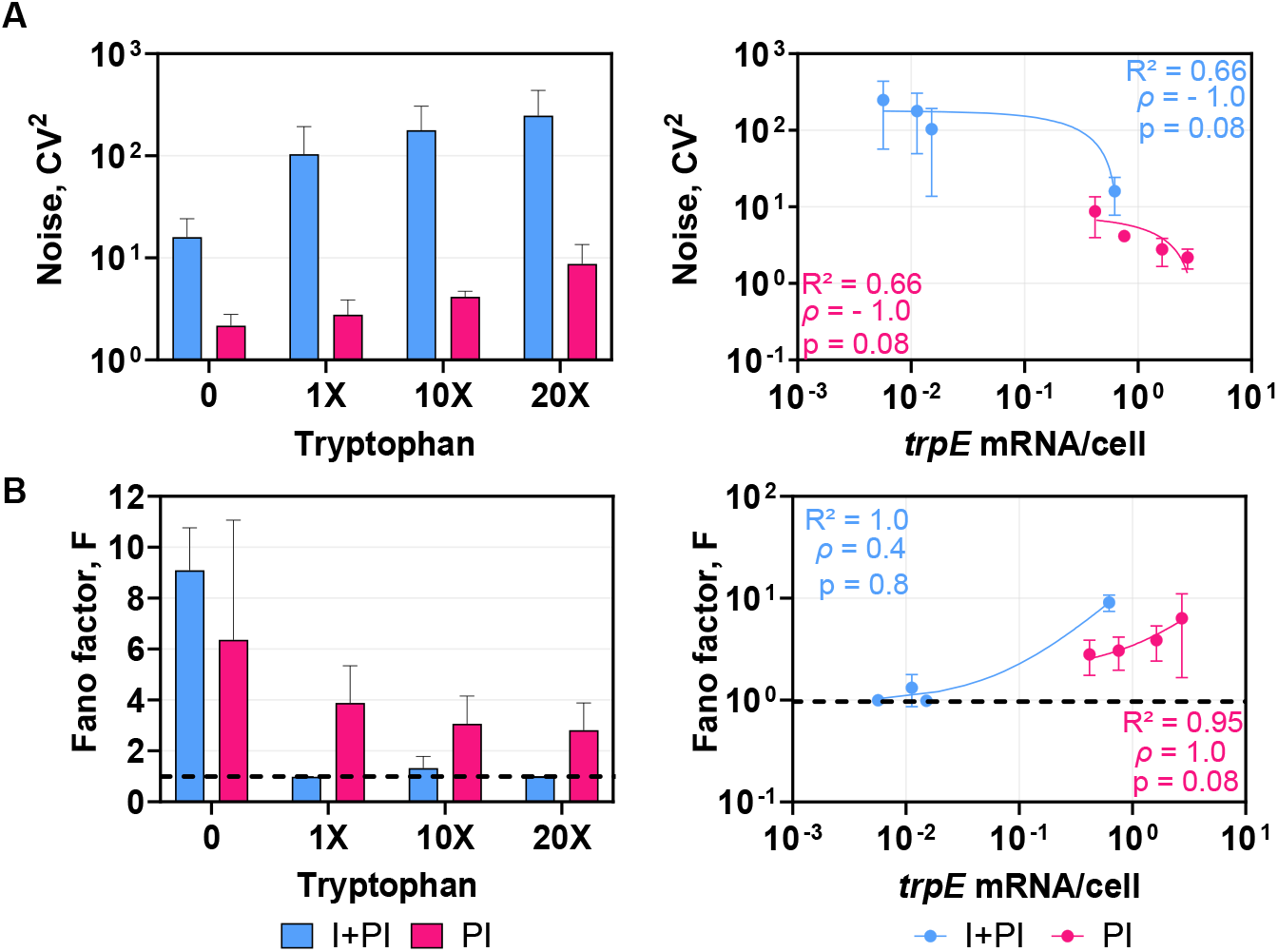
Comparison of noise and bursting of single- and dual-layer transcriptional regulation. (**A**) Noise (CV^2^ = σ^2^/μ^2^) and (**B**) burstiness (Fano factor F = σ^2^/μ) of *trpE* transcription under each tryptophan concentration (left) and as a function of the mean *trpE* mRNAs per cell (right). CV^2^ and F were calculated from the mean and standard deviations associated with the mRNA expression levels acquired from the data output from the Spätzcells software. The line at Fano = 1 distinguishes bursty event patterns (values above the line) from non-bursty patterns (values below the line). Linear regression lines were fitted to the right-hand plots with the corresponding R^2^ value displayed on each graph. Spearman’s rank correlation coefficient (*ρ*) and the associated p-values are also shown.

Further analysis revealed Fano factors >1 at 0X, in both modes of regulation, consistent with bursty transcription (Fig. 3B). In PI, burstiness declined gradually but persisted above 1 across all tryptophan levels. In I+PI however, burstiness dropped sharply with supplementation, approaching Poisson-like values (∼1), reflecting a switch to more uniform expression. Interestingly, *B. subtilis*, which lacks initiation control, also exhibited Fano factor >1, showing that burstiness originates downstream from TRAP-mediated attenuation rather than promoter activity.

Plotting Fano as a measure of burstiness against mean expression revealed a strong linear fit (R^2^ = 1 for I+PI; 0.95 for PI). However, correlation patterns differed: PI showed a strong monotonic relationship (*ρ* = 1, p = 0.08), consistent with gradual expression-dependent reduction in burstiness, whereas I+PI displayed a weaker monotonic trend (*ρ* = 0.4, p = 0.8) due to a plateau followed by an abrupt switch-like increase. Together, these results indicate that PI regulation produces a graded noise–burst response, while dual-layer I+PI regulation enforces abrupt, switch-like response to expression levels.

### Deconstruction of the PI regulation system

To determine the molecular regulator of transcription noise and burstiness in *B. subtilis* seen in Figure 3, we examined two central regulators: TRAP and its inhibitor anti-TRAP. To our knowledge, this is the first study assessing TRAP and anti-TRAP function at single-cell resolution (32,33,38,39). Figure 4 shows the effects of TRAP (Δ*mtrB*) and anti-TRAP (Δ*rtpA*) on transcriptional heterogeneity and bursting dynamics.

**Fig. 4.**
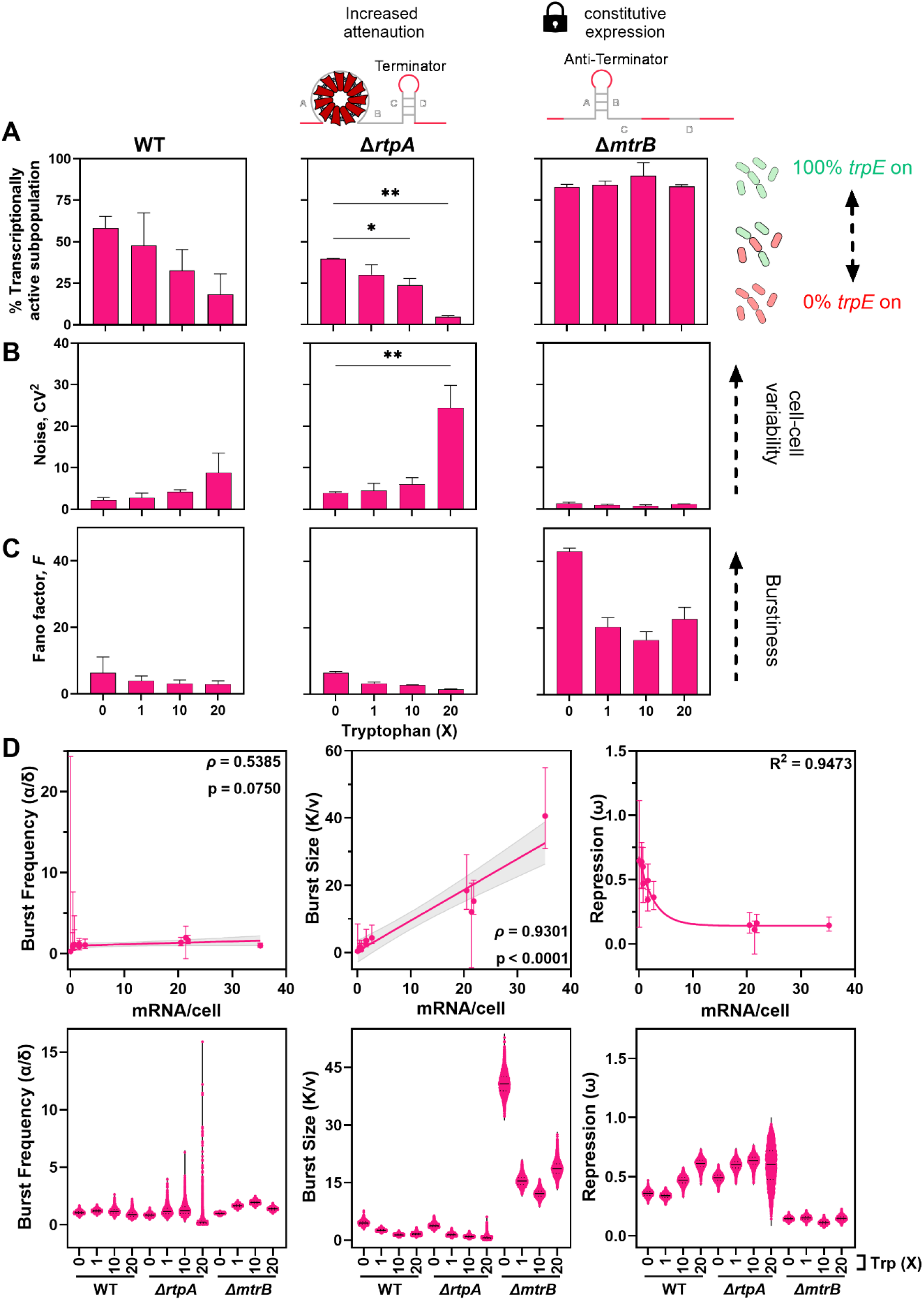
Transcriptional heterogeneity and bursting of regulation by attenuation only. Transcriptional heterogeneity was assessed in strains with either enhanced TRAP-mediated attenuation (Δ*rtpA*) or loss of TRAP-mediated attenuation (Δ*mtrB*), compared to the WT strain. The padlock symbol denotes a genetic “lock” state, in which Δ*mtrB* cells are transcriptionally fixed in the on state due to compromised attenuation regulation. Cells cultured with 0, 1X, 10X, or 20X tryptophan (0, 5, 50, and 100 µM). *trpE* expression was quantified using smRNA FISH. (**A**) the percentage of cells in the transcriptionally on state (defined as cells containing ≥1 *trpE* mRNA molecules). (**B**) Transcriptional noise (CV^2^ = σ^2^/μ^2^) and **(C)** Fano factor (F = σ^2^/μ) calculated from the mean (μ) and standard deviation (σ) of *trpE* mRNA copy number per cell. Mean values ± standard deviations are shown for two independent experiments. Statistical analysis was performed using one-way ANOVA followed by Dunnett’s multiple comparisons test (GraphPad Software). The dashed black line at Fano = 1 distinguishes bursty event patterns (values above the line) from non-bursty patterns (values below the line). (**D**) Burst kinetics of *trpE* transcription as a function of the mean *trpE* mRNAs per cell (top) and under each tryptophan condition tested (bottom).

In Δ*rtpA*, the fraction of transcriptionally active cells declined with increasing tryptophan, similar to WT, but significance was only reached at higher concentrations (10X, 20X) (Fig. 4A). This suggests anti-TRAP sustains a subpopulation of active cells under tryptophan-rich conditions, maintaining heterogeneity. By contrast, Δ*mtrB* showed >80% active cells across all conditions (Fig. 4A), indicating TRAP is required to establish and maintain a transcriptionally silent subpopulation in response to tryptophan. Noise patterns (Fig 4B) further supported this role. WT and Δ*rtpA* both showed increasing noise with tryptophan, implicating TRAP as a driver of cell-to-cell variability (Fig. 4B). However, Δ*rtpA* displayed higher noise than WT only at 20X tryptophan, consistent with anti-TRAP acting as a buffer against excessive TRAP-mediated repression under high tryptophan (Fig. 4B). In Δ*mtrB*, noise was consistently low, confirming TRAP is essential for introducing heterogeneity (Fig. 4B).

Bursting behaviour was broadly similar between WT and Δ*rtpA*, with moderate decreases in Fano factor at higher tryptophan but values always >1, confirming bursty transcription (Fig. 4C). By contrast, Δ*mtrB* showed consistently high burstiness under all tryptophan concentrations (Fig. 4C). Thus, TRAP suppresses bursts, while anti-TRAP exerts minimal effects on burst dynamics. We next used mathematical modelling (13) to dissect bursting parameters (Fig. 4D). In *B. subtilis*, repression (ω) was strongly inversely correlated with mean *trpE* expression (R^2^ > 0.9). In strains with functional TRAP (WT, Δ*rtpA*), repression increased with tryptophan, but anti-TRAP modulated sensitivity: Δ*rtpA* showed higher repression even in the absence of tryptophan, comparable to WT at 10X tryptophan. At 20X tryptophan, Δ*rtpA* repression was most heterogeneous, consistent with its elevated noise profile (Fig. 4.B). In Δ*mtrB*, repression remained low regardless of tryptophan, confirming TRAP as the main inducer of transcriptional silencing.

Burst size (Fig. 4D) positively correlated with mean expression (*ρ* = 0.93, p < 0.0001). WT and Δ*rtpA* showed modest tryptophan-dependent reductions, but removal of TRAP displayed a substantial increase, demonstrating that TRAP negatively regulates burst size. Burst frequency (Fig. 4D) correlated weakly and non-significantly with expression (*ρ* = 0.54, p = 0.08), suggesting lack of frequency regulation. TRAP deletion had no effect on frequency, though a small Δ*rtpA* subpopulation at 20X tryptophan showed high-frequency bursting, hinting that anti-TRAP may suppress rare cells with high transcriptional frequency states (Fig. 4D). Overall, TRAP-mediated attenuation primarily shapes bursting through control of burst size (Fig. 4D).

Taken together, these findings show that TRAP is essential for generating transcriptional heterogeneity and bursts by driving repression and limiting burst size, while anti-TRAP fine-tunes system sensitivity and buffers noise under high tryptophan.

### Deconstruction of the I+PI regulation system

We analysed *E. coli* regulatory mutants (Fig. 5) to dissect the contributions of transcriptional initiation (TrpR) and post-initiation regulation (attenuation). To isolate the contribution of ribosome-mediated attenuation, we used a Δ*trpR* strain lacking the TrpR repressor. In this strain, transcription is constitutively expressed at the point of initiation, and regulation of transcriptional output occurs through attenuation. To assess the effect of enhanced attenuation in the presence of regulation by TrpR we used a Δ*trpL* strain lacking the leader sequence necessary for ribosome stalling. In this strain, the absence of stalling leads to constitutive formation of the terminator hairpin, thereby genetically locking the system into a state of maximal attenuation. A double mutant (Δ*trpR*Δ*trpL*) lacking both TrpR and the leader sequence was used to investigate the effect of maximal attenuation in the absence of initiation regulation. Although recent studies have identified small RNAs that interact with the *trpL* 5’ UTR in *E. coli* and may influence *trpE* expression, our primary emphasis is on repression occurring at the initiation stage by TrpR and ribosome-mediated attenuation (40).

**Fig. 5.**
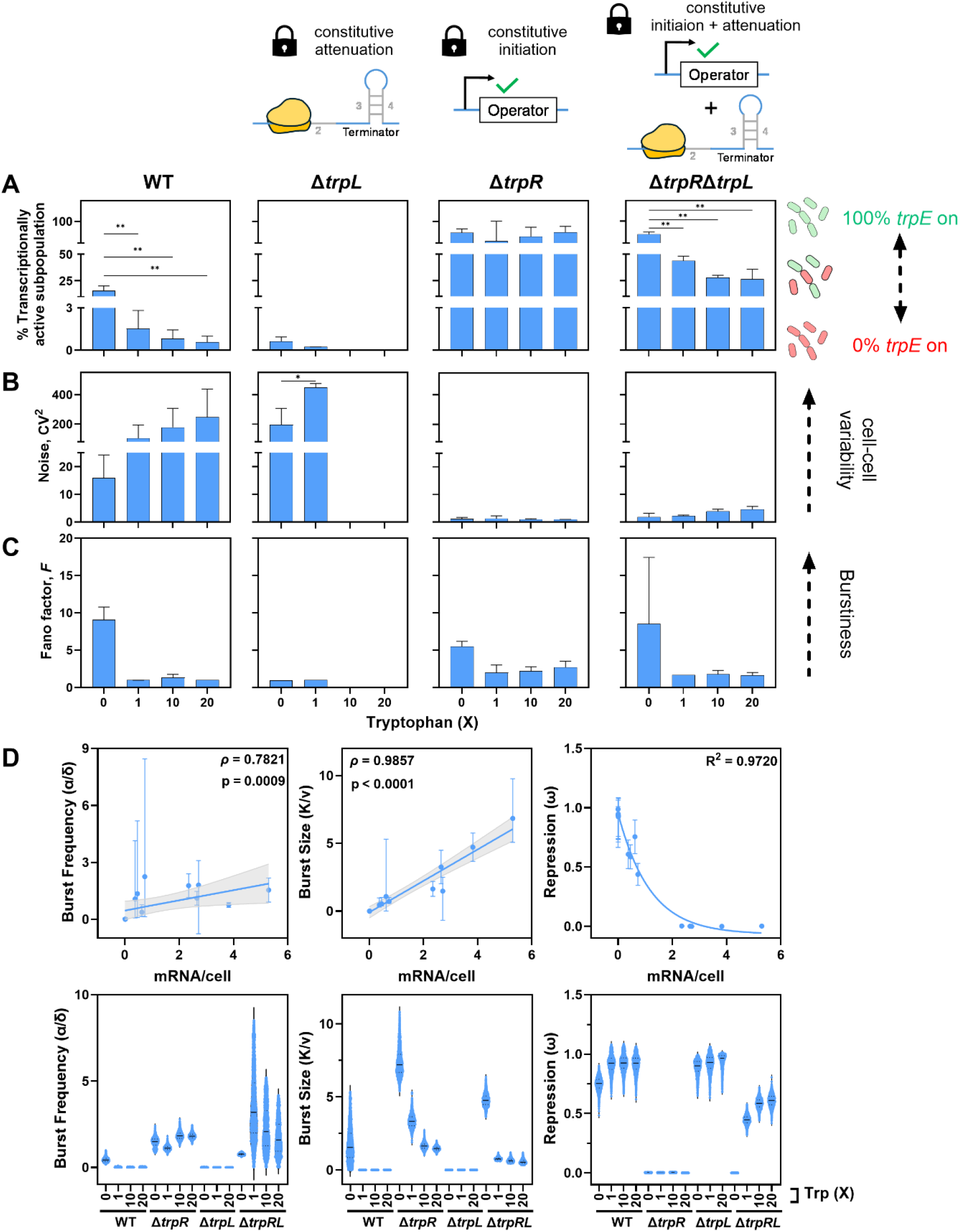
Transcriptional heterogeneity and bursting of combined regulation by repression and attenuation. Transcriptional heterogeneity was assessed in strains with enhanced ribosome mediated attenuation (Δ*trpL*), constitutive initiation (Δ*trpR*) or both (Δ*trpR*Δ*trpL*), compared to the WT strain with functional initiation and post-initiation regulation. The padlock symbol denotes a genetic “lock” state, in which cells are either genetically fixed in the on state for transcription attenuation (Δ*trpL*), initiation (Δ*trpR*) or both (Δ*trpR*Δ*trpL*). Cells cultured with 0, 1X, 10X, or 20X tryptophan (0, 5, 50, and 100 µM). *trpE* expression was quantified using smRNA FISH. (**A**) The percentage of cells in the transcriptionally on state (defined as cells containing ≥1 *trpE* mRNA molecules). (**B**) Transcriptional noise (CV^2^ = σ^2^/μ^2^) and **(C)** Fano factor (F = σ^2^/μ) calculated from the mean (μ) and standard deviation (σ) of *trpE* mRNA copy number per cell. Mean values ± standard deviations are shown for two independent experiments. Statistical analysis was performed using one-way ANOVA followed by Dunnett’s multiple comparisons test (GraphPad Software). (**D**) Burst kinetics of *trpE* transcription as a function of the mean *trpE* mRNAs per cell (top) and under each tryptophan condition tested (bottom).

In Δ*trpR*, ∼70% of cells remained active across all tryptophan concentrations, showing attenuation alone cannot silence large subpopulations (Fig. 5A). The mutant Δ*trpL* showed decreasing activity with tryptophan, reaching full repression at ≥10X. Δ*trpRΔtrpL* resembled Δ*trpR* in the absence of tryptophan (∼79% active) but declined with supplementation (Fig. 5A). Thus, TrpR primarily dictates population transcriptional activity, with influence by attenuation only when maximised.

Analysis of transcriptional noise (Fig. 5B) revealed Δ*trpL* exhibited elevated noise without tryptophan compared to the WT, peaking at 1X, but dropped to zero when fully repressed (10– 20X) (Fig. S4). Δ*trpR* maintained low noise regardless of tryptophan, while Δ*trpR*Δ*trpL* showed moderate increase of noise at high tryptophan (Fig. 5B). Strains lacking TrpR consistently had lower noise, implicating TrpR as the main source of variability, with limited impact from genetically maximised attenuation (Fig. 5B).

Bursting analysis (Fig. 5C) showed that Δ*trpL* is non-bursty at 0 and 1X tryptophan. By contrast, Δ*trpR* and Δ*trpRΔtrpL* remained bursty (Fano factor > 1) under all conditions, indicating that absence of TrpR restores bursty transcription even with maximal attenuation.

Repression (Fig. 5D) inversely correlated with mean *trpE* expression (R^2^ > 0.9). WT and Δ*trpL* showed high repression with tryptophan, but only Δ*trpL* was elevated without it, consistent with attenuation being relieved only under starvation (29). Δ*trpR* showed minimal repression, while Δ*trpRΔtrpL* increased repression with tryptophan (Fig. 5D).

Burst size (Fig. 5D) strongly correlated with mean expression (*ρ* = 0.99, *p* < 0.0001) and declined in WT and Δ*trpR* with tryptophan supplementation, however Δ*trpR* retained leaky bursts (Fig. 5D). Δ*trpRΔtrpL* further reduced burst size with tryptophan, highlighting post-initiation effects (Fig. 5D). Δ*trpL* remained low, indicating complete repression of burst size requires both initiation and attenuation (Fig. 5D).

In addition, burst frequency (Fig. 5D) also correlated with expression (*ρ* = 0.78, *p* = 0.0009). WT frequency decreased with tryptophan while Δ*trpR* remained high. Δ*trpL* was reduced, and Δ*trpRΔtrpL* declined with supplementation (Fig. 5D). Together, these data show that transcription of *trpE* is controlled through coordinated regulation of both burst size and frequency. To our knowledge, this is the first report of a σ^70^-dependent promoter capable of regulating both the frequency and size of transcriptional bursts.

## Discussion

Variability in mRNA levels arises from transcriptional bursting, an inherently stochastic process producing intermittent bursts of transcripts (3,5–8,10). These fluctuations are a major source of transcriptional noise in both eukaryotes and prokaryotes (*2,14,16,18,23*). Most work has focused on transcription initiation, where bursts are modulated either by frequency or burst size, depending on promoter architecture (*2,13–19*). Early models proposed uniform bursting in bacteria (*44*), but growing evidence shows that gene-specific features, including sigma factors, modulate bursting dynamics (*2,13–15)*. In contrast, how post-initiation mechanisms such as premature termination shape bursting remains poorly understood.

Here we demonstrate for the first time that enhancer-independent transcription can also be regulated by burst frequency through the combined action of initiation and attenuation control. Exploring such regulatory architectures is key to understanding bacterial strategies for promoting heterogeneity and stress survival. These insights also advance synthetic biology, enabling the design of circuits that either minimise variability or exploit it for desired outcomes.

We further show that, at the native locus of biosynthetic pathways, transcriptional states in one subpopulation can influence others via metabolic cooperation, revealing a previously unrecognised source of heterogeneity through metabolite exchange. This dynamic not only reduces the metabolic burden and creates functional heterogeneity within the population but also stabilises cooperative interactions and shapes overall community structure.

Comparing I+PI versus PI-only regulation, we observed the canonical inverse relationship between noise and mean expression associated with σ^70^-dependent genes and found that initiation control introduces additional cell-to-cell variability by adding a regulatory “decision point.” Dissection of the *trp* operon shows that in *B. subtilis*, TRAP-mediated attenuation is a key regulator of noise, whereas in *E. coli*, both TrpR and ribosome-mediated attenuation act as molecular switches. This reveals that even in promoters regulated at initiation, downstream attenuation can influence cell-to-cell variation.

The differing regulatory architectures give rise to distinct patterns of transcriptional bursting, which may reflect contrasting adaptive strategies. In the PI-only system of *B. subtilis*, TRAP-mediated attenuation modulates burst size but not frequency, preserving heterogeneity. This strategy is consistent with maintaining subpopulations of high *trpE* expression, and potentially advantageous in its natural habitat. Soil is a spatially structured environment, where nutrient availability can vary dramatically even over micrometer scales. Subpopulations of *B. subtilis* with high *trpE* expression could locally produce tryptophan, support nearby cells and potentially enable metabolic cooperation. This could help maintain community stability and support growth in micro-niches within the soil where tryptophan is scarce. By contrast, the I+PI system of *E. coli* allows regulation of both burst size and frequency. This suppresses *trpE* expression efficiently while providing temporal flexibility for rapid responses to the dynamic environment in the gut where *E. coli* faces episodic scarcity and abundance of nutrients.

Our study shows how transcriptional noise and bursting in bacteria arise from the interplay between transcription initiation and post-initiation regulation. Building on the established view that transcriptional bursting is dictated by initiation, we show that premature termination of transcription can influences both burst frequency and size. Moreover, we highlight the previously underappreciated role of metabolic cross-feeding in driving transcription heterogeneity. Overall, our work underscores the complexity of transcriptional regulation beyond initiation events, revealing new molecular switches that shape gene expression variability and highlighting the need to consider not only genetic circuitry, but also community and environmental context to fully understand and engineer microbial behaviour.

## Materials and Methods

### Bacterial strains

The *E. coli* BW25113 (WT) and Δ*trpR*, Δ*trpE* and Δ*trpL* single-gene knockouts were acquired from the Keio collection (all in the BW25113 background) (41). Mutant stains were rendered markerless by removing the kanamycin resistance cassette situated at the native locus of the target gene, using FLP recombinase from the pCP20 plasmid. The Δ*trpR*Δ*trpL* double mutant, was generated by P1 phage transduction using the Δ*trpL* strain as the donor and the Δ*trpR* strain as the recipient. *T*he Δ*trpE*, Δ*mtrB*, Δ*rtpA* knockouts of *B. subtilis* were obtained from the *Bacillus* Genetic Stock Centre (BGSC) in a tryptophan auxotrophic (*trpC*2) background (42). Kanamycin markers of each mutant acquired was PCR amplified; gel purified and transformed into tryptophan prototrophic WT (168 *trp*C+) using natural competence. Briefly, a single colony of donor *B. subtilis* tryptophan prototrophic WT (168 *trp*C+) strain was inoculated into SP medium at 37 °C with shaking (150 rpm) O/N. Then diluted 1:50 with 500 µL of fresh SP medium and 1 µL of purified PCR product and incubated at 37 °C with shaking (150 rpm) for 5.5 hours. Cultures were then centrifuged and resuspended in 1 mL of LB medium and incubated for 1.5 hours at 37 °C with shaking (150 rpm). Cultures were then plated on LB medium supplemented with 5 µg/mL kanamycin. Kanamycin markers were excised using pDR244, Cre recombinase expressing plasmid, obtained from the BGSC (42). The Δ*mtrB* strain was transformed with the SG13 plasmid, containing a *Pveg-gfp* transcriptional fusion, acquired from BGSC (42) using natural competence. The resultant Δ*mtrB amyE::gfp* was used in the smRNA FISH co-culture experiment.

### Growth assay

A single colony was inoculated into 5 mL of LB broth and incubated at 37 °C, with shaking at 220 rpm. The next day, cell cultures were centrifuged at 5000 rpm for 5 minutes and supernatant was removed. Pellet was washed thrice in 1.5 mL of medium of interest (fresh M9 minimal media) each time centrifuging at 5000 rpm for 5 minutes. The OD600 was standardised to 0.1 in 500 μL. A volume of 100 μL of the experimental culture was transferred into a flat bottom well of a 96-well microtiter plate, including three replicates of each condition. The OD600 was measured every hour for 40 hours using a FLUOstar® Omega (BMG LABTECH) UV/vis filter-based microplate reader

### Single-molecule RNA FISH

Fluorescently labelled probes targeting the *trpE* mRNA transcripts were designed for both *E. coli* and *B. subtilis* organisms using Stellaris® Probe Designer software (LGC Biosearch Technologies). Design parameters included an oligonucleotide length of 20 nucleotides, a minimum spacing of 2 nucleotides between probes, and a masking level set to 1–2. All probes were conjugated with 6-carboxytetramethylrhodamine succinimidyl ester (6-TAMRA) as the fluorescent dye. A single colony of the target strain was picked using a sterile loop and transferred into 5 mL of LB broth in a sterile 30 mL polystyrene universal tube and incubated at 37 °C, with shaking at 220 rpm. The next day, cell cultures were centrifuged at 5000 rpm for 5 minutes and supernatant was removed. Pellet was washed thrice in 1.5 mL of fresh modified M9 medium each time centrifuging at 5000 rpm for 5 minutes. The OD600 was standardised to 0.1 in 20 mL culture volume supplemented with 0, 5, 50 or 100 µM (corresponding to 0, 1, 10 and 20X) tryptophan, and harvested at mid-exponential phase (OD600 0.4) by centrifugation. The cells were resuspended and fixed in 1 ml of ice-cold 1X PBS in DEPC-treated water containing 3.7% (v/v) formaldehyde and incubated for 30 minutes at room temperature. After fixation, the cells were washed twice with 1 ml of 1X PBS in DEPC-treated water, then resuspended in 1 ml of 70% (v/v) ethanol in DEPC-treated water and incubated for 1 hour at room temperature to permeabilise. Following permeabilisation, the cells were washed with 1 ml of 2X SSC in DEPC-treated water containing 40% (w/v) formamide and incubated overnight at 30°C with hybridisation buffer (2X SSC in DEPC-treated water, 40% (w/v) formamide, 10% (w/v) dextran sulfate, 2 mM ribonucleoside-vanadyl complex, and 1 mg/ml *E. coli* tRNA) and 1 μM *trpE* specific fluorescent probes. After hybridisation, 10 μl of the cells were washed twice in 200 μl of ice-cold wash solution (40% (w/v) formamide and 2X SSC in DEPC-treated water) and incubated for 30 minutes at 30°C. The chromosomal DNA was then stained with DAPI-containing wash solution (40% (w/v) formamide, 2X SSC in DEPC-treated water, and 10 µg/ml DAPI) for 30 minutes at 30°C. The cells were resuspended in 100 μl of 2X SSC in DEPC-treated water, from which 2 μl was spotted onto the centre of a 1% (w/v) agarose gel. Once dry, a 1x1 cm square was cut around the sample and transferred to a 76 x 26 mm microscope glass slide.

### Quantification of *trpE* mRNA molecules

Cells were imaged using a Leica Stellaris 8 confocal microscope, acquiring five z-slices at 200 nm intervals for each channel (brightfield, DAPI and 6-TAMRA) across multiple x/y (stage) positions. The resulting 16-bit .tif images from all three channels were used to generate cell segmentations using the Schnitzcells software (43) in MATLAB (MathWorks). The *trpE* mRNA copy numbers in single cells were quantified with the Spätzcells program (37) in MATLAB, using the 6-TAMRA channel images and the cell segmentations. Fluorescent spots within the segmented cells were detected and differentiated from nonspecific background signals by setting a false-positive threshold using *ΔtrpE* cells as a negative control. False-positive spots were excluded by setting the threshold at the 99.9th percentile of spot intensities observed in *ΔtrpE* cells. Fluorescent spots exceeding the false-positive threshold were classified as specific signals corresponding to *trpE* mRNA molecules hybridised with complementary DNA probes. These fluorescent spots’ peak height and intensity were analysed to determine the mRNA copy numbers. The intensity distribution of spots from a low-expressing control strain was fitted to a multi-Gaussian function, with the mean of the first Gaussian representing the intensity of a single mRNA molecule. The total fluorescence intensity of spots in each cell was divided by the intensity of a single mRNA molecule to calculate the number of mRNA molecules per cell. These data were then used to calculate the relative frequencies, mean and standard deviation of mRNA copy numbers across the population.

### Co-culture smRNA FISH

Single colonies of WT and Δ*mtrB* were inoculated in 5 mL of LB broth, in separate tubes and incubated at 37 °C, with shaking at 220 rpm O/N. The cultures were then centrifuged at 5000 rpm for 5 minutes and supernatant was removed. The pellet was washed thrice in 1.5 mL of modified M9 medium each time centrifuging at 5000 rpm for 5 minutes. The OD600 of each strain was standardised to 0.05 in a 20 mL culture volume of fresh modified M9 medium and incubated at 37 °C, with shaking at 220 and harvested at mid exponential phase (OD600 0.4) by centrifugation. Once harvested the cells were treated for smRNA FISH as previously described. Importantly for the co-culture experiment, along with brightfield, DAPI, and 6-TAMRA channel images, GFP images were also captured to distinguish the GFP fluorescently tagged Δ*mtrB* cells within the images such that only cell segmentation of non GFP fluorescent WT cells were generated using the Schnitzcells software in MATLAB (MathWorks).

### Amino acid measurements

Single colonies of *E. coli and B. subtills* WT strains (BW25113 and 168 *trpC+*, respectively) were grown overnight in 5 mL of LB broth at 37 °C, with shaking at 220 rpm. The next day the cultures were centrifuged at 5000 rpm for 5 minutes and supernatant was discarded. The pellets were then resuspended and washed in M9 minimal medium thrice, each time centrifuging at 5000 rpm for 5 minutes. The OD600 was adjusted to 0.1 in 15 mL of M9 minimal media and incubated at 37 °C, with shaking at 220 rpm for 24 hours. 1 mL of the culture was filter sterilised with Millipore 0.2 µm filter to remove the cells. The spent media was then immediately frozen in liquid nitrogen and stored at -80 C. The amino acid from the spent medium were derivatised using the AccQ-Tag kit (Waters) as per the manufacturer’s guidelines and quantified by tandem mass spectrometry using a TQSµ coupled to a Acquity UPLC equipped with HSS T3 2.1 × 150 mm, 1.8 μm column, i. Separation was achieved using a gradient with phase A water with 0.1% formic acid (v/v) and acetonitrile with 0.1% formic acid (B) and the column held at 45 C. Gradient elution was performed with 4% B at 0.6 mL/min starting, held for 0.5 min, then to 10% B over 2 min, then to 28% B over 2.5 min and to 95% B for 1 min, before returning to 4% B (1.3 min) for re-equilibration. The amino acid derivatives were quantified in positive modeas published previously (44). Data were analysed using an in-house Matlab pipeline based on software published by Behrends et al. (2011) (45). Values are expressed as arbitrary units (A/U).

### Computational Analysis of burst kinetics

Burst size, frequency, and transcriptional repression were estimated using a Bayesian inference framework as described in (13). A zero-inflated negative binomial model was used to describe the distribution of mRNA copy numbers. Parameters θ=[ω,r,p] were inferred using Metropolis-Hastings MCMC sampling, with the posterior distribution being computed by the product of the likelihood and the priors. Uniform priors were used for ω and p (0,1), while r was assigned from a half-normal (µ=0, σ=20) positively truncated prior. Sampling was performed using a custom MCMC implementation with a multivariate Gaussian proposal distribution, in which the standard deviation was set to 5% of the current parameter values, to ensure good convergence. Chains were iterated for 500,000 steps, with 100,000 discarded as burn-in and thinning applied by a factor of 100. Burst parameters, maximum a posteriori estimate (measure of centre of the error bar) and 95% credible intervals were computed from the posterior distributions for further interpretation.

## Supporting information

Supplementary information Moradian, Ali et al

## Acknowledgments

We would like to thank Nils Averesch for providing us with the tryptophan prototrophic 168 strain of *B. subtilis*. We would also like to thank Michaela Egertova for microscopy support. This work was supported by grants from the Leverhulme Trust (RPG-2021-050) and the BBSRC (BB/W019698/1).

