## Supplementary information Moradian, Ali et al for "Single-cell Analysis of Attenuation-Driven Transcription Reveals New Principles of Bacterial Gene Regulation"

Supplementary Information for Moradian, Ali et al

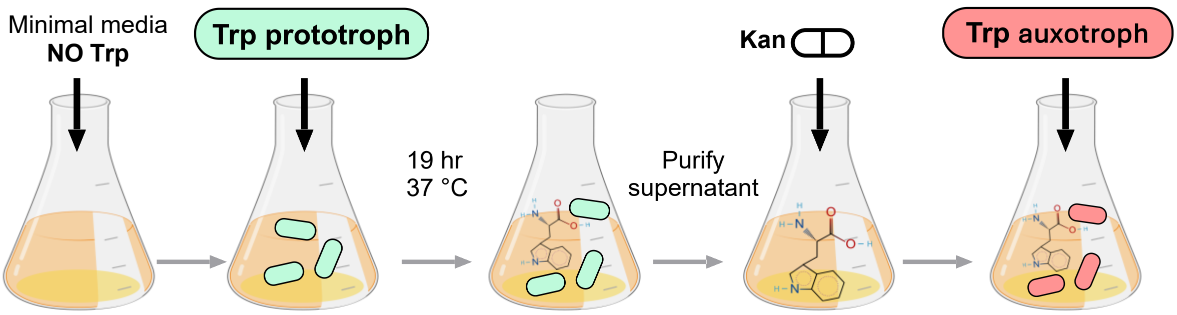


**Fig. S1. Experimental set up using tryptophan auxotroph's as a biosensor.** Tryptophan prototrophs were cultured in minimal media lacking tryptophan. Following the purification of the supernatant after a 19-hour incubation, kanamycin was added and the tryptophan auxotroph strain (Δ*trpE*::*kan*) was cultured and growth was monitored. Growth data is presented in supplementary Fig. S2.

| Amino acid | *B. subtilis* | *E. coli* |
|---|---|---|
| Glycine | Present | Present |
| Alanine | Present | Present |
| Leu/Ile | Present | Present |
| Glutamate | Present | Present |
| Phenylalanine | Present | Present |
| Tyrosine | Present | Present |
| Tryptophan | Present | Present |
| Lysine | Absent | Absent |
| Asparagine | Absent | Absent |
| Aspartate | Absent | Absent |
| Methionine | Absent | Absent |
| Histidine | Absent | Absent |
| Arginine | Absent | Absent |
| Citrulline | Absent | Absent |

**Table S1. Table of amino acids identified either as present or absent in the supernatant of WT cells at 24 hours post inoculation using mass spectrometry.**

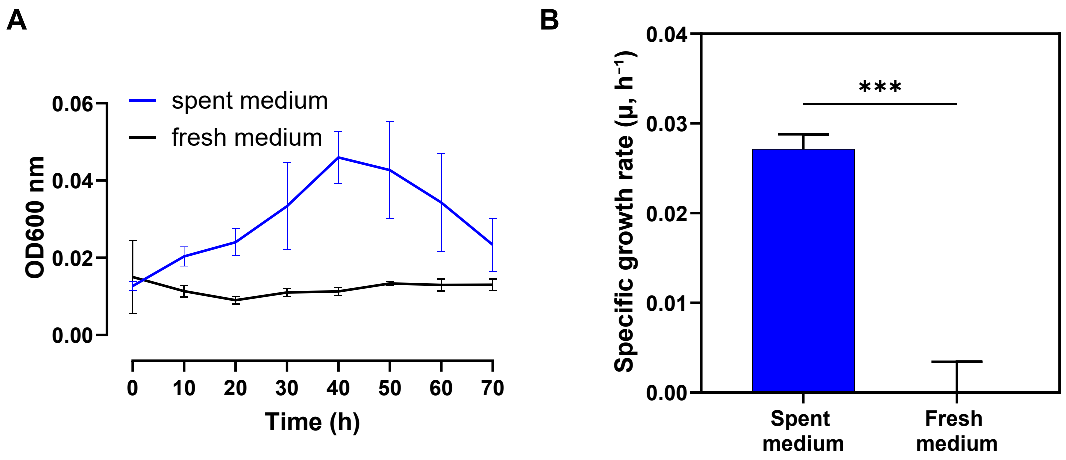


**Fig. S2. Tryptophan cross feeding between tryptophan prototrophic donners and auxotrophic recipients.** (A) Growth curve of a *B. subtilis* tryptophan auxotroph strain (Δ*trpE*) cultured in either fresh minimal medium or spent medium collected from tryptophan prototrophic WT. Bacterial growth was monitored over 10 hours over 70-hour OD600 using an Omega microplate reader in 100 μL culture volumes. Data represent the mean ± standard deviation from three independent experiments. (B) Specific growth rate (μ, $h^{-1}$) of the tryptophan auxotroph cultured in spent versus fresh medium, calculated from OD600 values between 10 and 40 hours using the exponential growth rate formula (μ = ln(ODt2)−ln(ODt1) / t2−t1). Bars represent the mean ± standard deviation of three biological replicates. Statistical significance was determined using an unpaired *t*-test (p = 0.0002) (GraphPad Software Inc.).

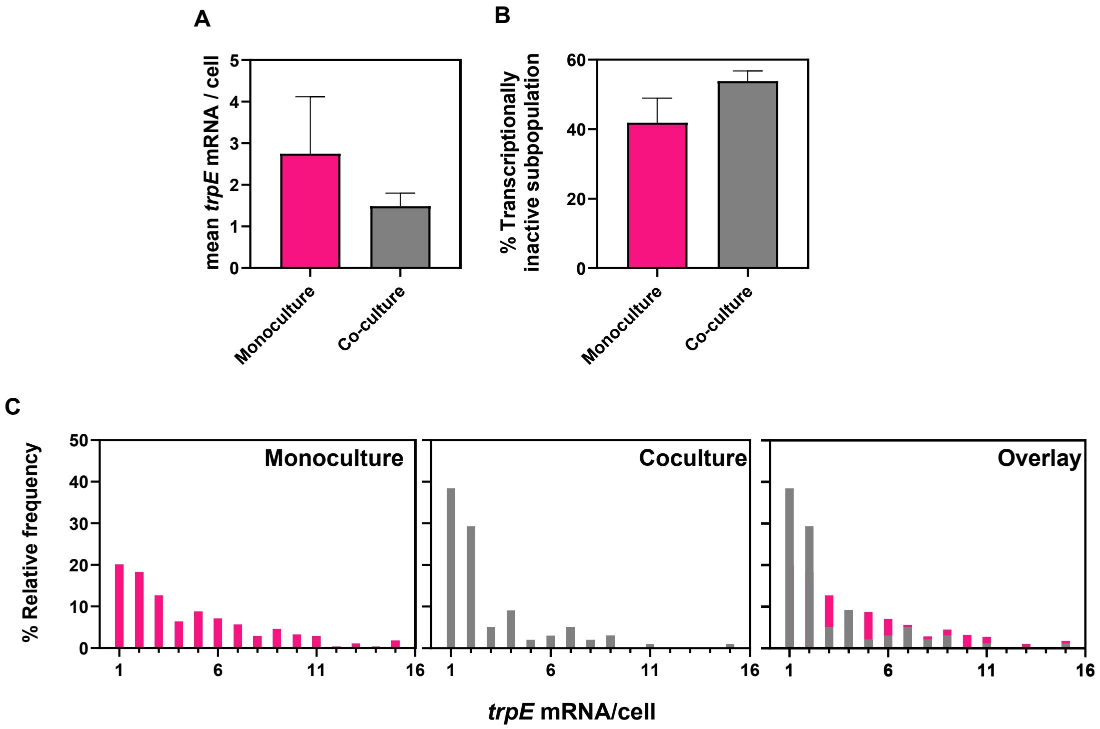


**Fig. S3. Co-culture with high *trpE*-expressing mutant suppresses *trpE* transcription dynamics in wildtype cells.** Expression dynamics of *trpE* in WT cells grown either as a monoculture or in co-culture with a high *trpE*-expressing TRAP mutant (Δ*mtrB*). Co-cultures were established at a 1:1 ratio of WT and Δ*mtrB* cells in minimal medium without supplemented tryptophan, grown to mid-exponential phase, and fixed with formaldehyde prior to hybridisation with TAMRA-labelled DNA probes targeting *trpE* mRNA. Cells were imaged using a Leica Stellaris 8 confocal microscope. Using Schnitzcells cell segmentation software (43), WT cells lacking GFP fluorescence were selected for quantification of *trpE* mRNA. (A) The mean number of *trpE* mRNA molecules per WT cell, quantified using smRNA FISH (37), in the WT monoculture and WT + Δ*mtrB* co-culture condition. (B) The proportion of WT cells in a transcriptionally inactive state (defined as having ≥1 *trpE* mRNA molecule), in the WT monoculture and WT + Δ*mtrB* co-culture condition. (C) Percentage relative frequency of *trpE* mRNA copy numbers in transcriptionally active (defined as cells expressing 1 or more molecules of *trpE* mRNA) WT cells in the WT monoculture and WT + Δ*mtrB* co-culture condition.

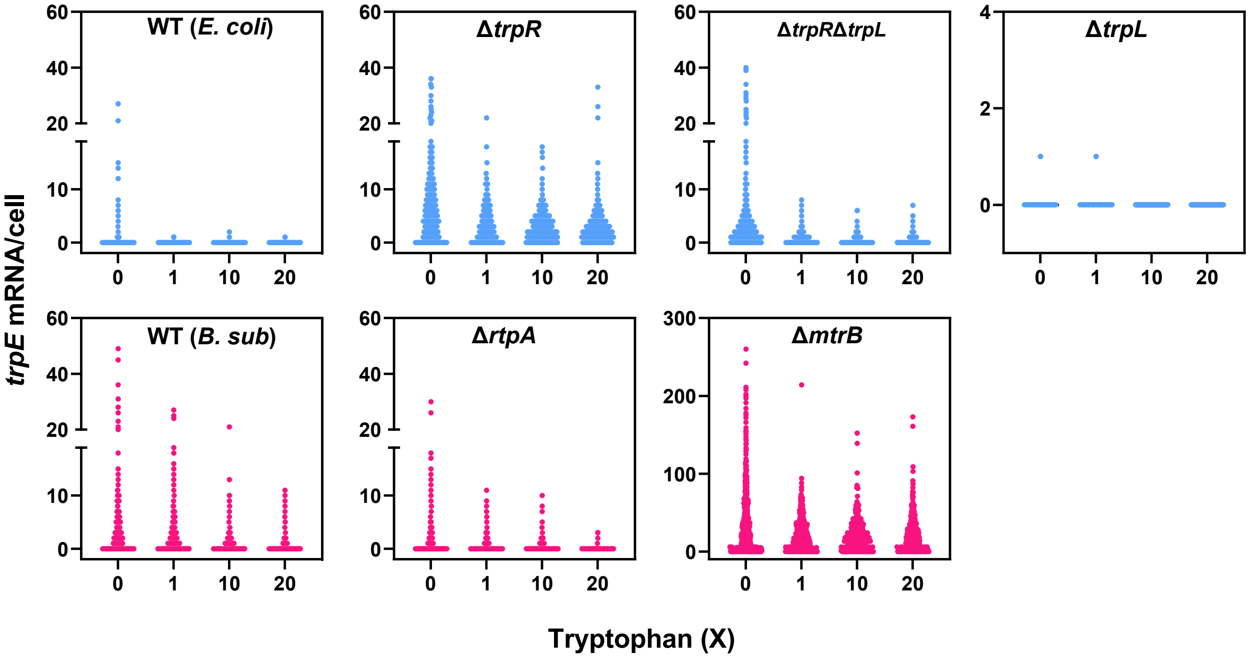


**Fig. S4. Single molecules of *trpE* mRNA quantified in single cells of *B. subtilis* and *E. coli* strains in response to environmental tryptophan.** Single molecules of *trpE* mRNA were quantified using Spätzcells software (37), and the resulting mRNA distributions are shown as scatterplots. The top panel (blue) represents all *E. coli* strains analysed in this study, while the bottom panel (pink) shows all *B. subtilis* strains.

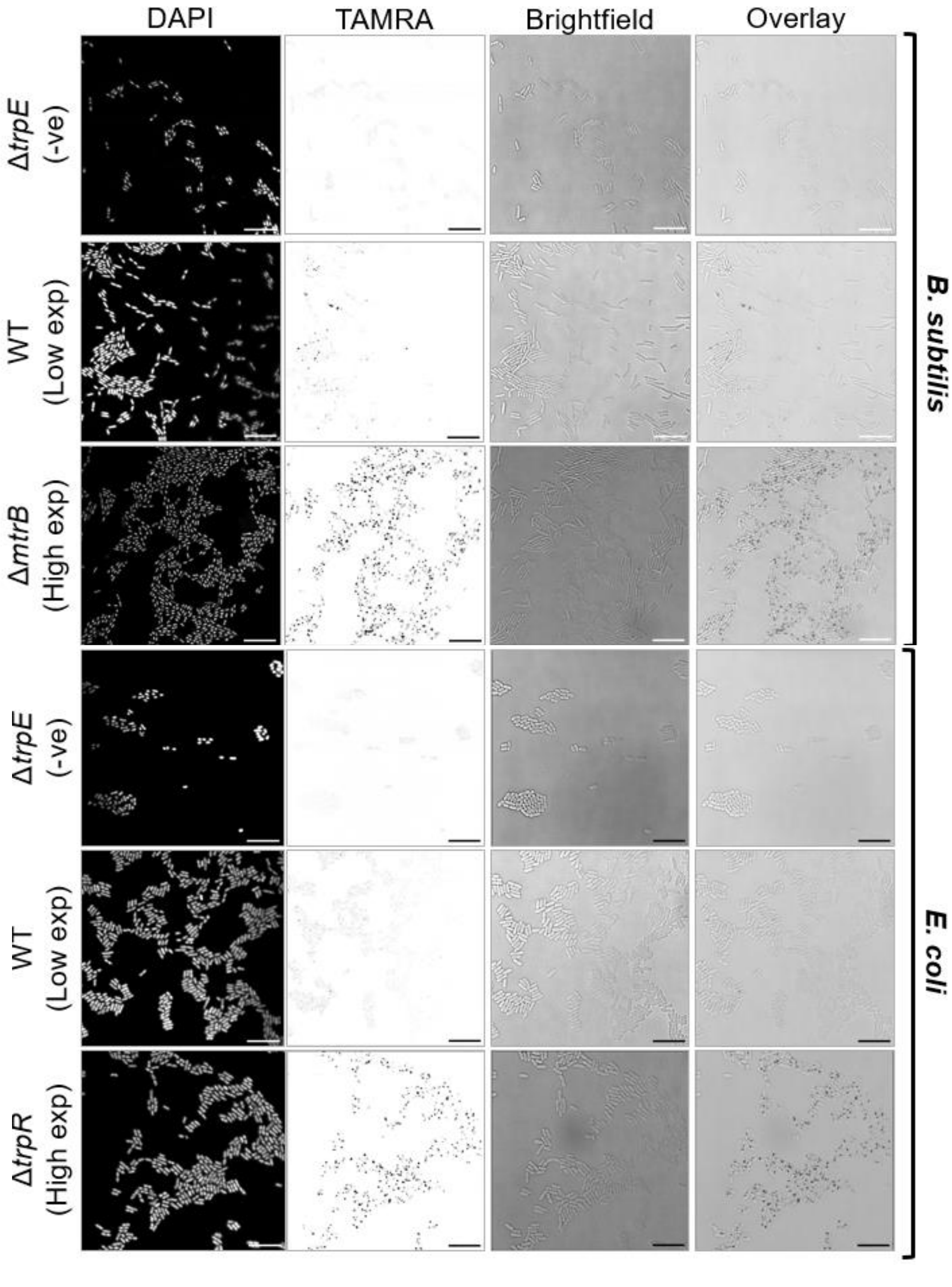


**Fig. S5. Representative smFISH images targeting *trpE* in *B. subtilis* and *E. coli*.** Shown are example images of DAPI, TAMRA, brightfield, and TAMRA + brightfield overlays for *E. coli* and *B. subtilis*. The Δ*trpE* negative controls for both species display an absence of TAMRA signal. WT cells of both species show low TAMRA signal. The Δ*mtrB B. subtilis* mutant, which constitutively expresses *trpE* at both transcription initiation and post-initiation, exhibits a markedly increased TAMRA signal relative to the WT. Similarly, the Δ*trpR E. coli* mutant, which constitutively initiates *trpE* transcription, also shows increased TAMRA signal compared to the WT.

| *E. coli* | | | | |
|---|---|---|---|---|
| | **WT** | | | |
| **Tryptophan (X)** | **0** | **1** | **10** | **20** |
| **Mean** | 0.623 | 0.015 | 0.011 | 0.006 |
| **SD** | 0.152 | 0.009 | 0.008 | 0.003 |
| | **Δ*trpR*** | | | |
| **Tryptophan (X)** | **0** | **1** | **10** | **20** |
| **Mean** | 5.298 | 2.654 | 2.345 | 2.710 |
| **SD** | 2.173 | 2.014 | 0.856 | 0.629 |
| | **Δ*trpL*** | | | |
| **Tryptophan (X)** | **0** | **1** | **10** | **20** |
| **Mean** | 0.006 | 0.002 | 0 | 0 |
| **SD** | 0.002 | 0.00009 | 0 | 0 |
| | **Δ*trpR* Δ*trpL*** | | | |
| **Tryptophan (X)** | **0** | **1** | **10** | **20** |
| **Mean** | 3.823 | 0.731 | 0.460 | 0.378 |
| **SD** | 1.431 | 0.045 | 0.024 | 0.122 |
| ***B. subtilis*** | | | | |
| | **WT** | | | |
| **Tryptophan (X)** | **0** | **1** | **10** | **20** |
| **Mean** | 2.731 | 1.632 | 0.758 | 0.418 |
| **SD** | 0.971 | 0.831 | 0.256 | 0.249 |
| | **Δ*rtpA*** | | | |
| **Tryptophan (X)** | **0** | **1** | **10** | **20** |
| **Mean** | 1.663 | 0.750 | 0.487 | 0.059 |
| **SD** | 0.029 | 0.131 | 0.087 | 0.003 |
| | **Δ*mtrB*** | | | |
| **Tryptophan (X)** | **0** | **1** | **10** | **20** |
| **Mean** | 35.214 | 21.814 | 21.411 | 20.495 |
| **SD** | 6.568 | 1.443 | 1.795 | 3.631 |

**Table S2. The mean and standard deviation of *trpE* mRNA levels quantified in every condition and strain tested in this study using smRNA FISH.**

| A | *E. coli* | |
|---|---|---|
| tttgtgtttttggctgagag | ccttttattgacgggactaa | caagtgacctgtctctagag |
| gaatagcgctgttagggtgg | aatagagcgactttgcgact | cggcataacttgacctttac |
| aaaagtggtcaacacacccc | actggtagtctttttttcgt | atggctagtatttctcgaca |
| cttaggcgtctatagctgtc | cataagtccggtcggacaaa | cttgtagactacgaccaact |
| actaaatttttcggacgacg | ctttttgttgcagagtgacg | cattactagaccgtgcgtaa |
| cgacgcgtaatgtcgaaatc | gacttgcttgatgcagtcgt | ggctagagtggtttcaactg |
| cagtgttaggtccgtgaaag | ggcgtatacgcaacacttac | aaggatacactacgtggagc |
| gaggaccgtgatgacctatt | tagtctcgctacttctcaag | cttgacgcagtgctagaact |
| tcacttgttagtggtttgac | acgcaaacaacgtttttcgc | atacttataccccctgcaatt |
| gacagtcaggtgacgaccta | cgacctctttaaaaggtcca | gcgatacgtcaattaacggc |
| gcgaatacgagggaaagcca | tagagcggcaaagagagacg | catccaataaagtggcgcgt |
| actgcgaaaggcaaataacg | ccggataatgcacgactttt | ctagagctgtggacgtaaca |
| gacaacttacatggcttcct | agggtcgggcatgtacaaaa | acgaccacatcaggaactaa |
| tgctcttcggtacaagaagc | taaagtgggataaaccgcgc | gctgctttgggcattgtttc |
| ccggacaagagaatactgga | ctttcgagcgagttcatact | gcgacatgacgcgcgataac |
| cgccctaaacttctaaatgg | ggtctaactctagatgggct | gtagtacgtgtcctctgaaa |

| B | *B. subtilis* | |
|---|---|---|
| ggcgtaaaaatctcctgtcg | cggcctgtaattaacgcata | gcctcgtaatgtacgagcaa |
| gctaacacctctggaagtgt | tactttggtttttgcaggtg | agaacgggctttgctatagc |
| gctatgtgactgtgggtaag | ataggttatacgttccgagt | tcatcgtctcataccaagac |
| tactatctcttcgaactgtc | tctcctttgttttttgcttt | cggcctcaagtgtttttaac |
| agaagaactttcgttcctga | ggtagttttagtagacctcg | acgtgtaatagagccaccaa |
| tgtaggtgaaccaggtctat | tttactacctggtttttttg | gctaacttttttccccaagt |
| gcaaatagccggacttaggt | tatgttctgtgggtcgaaac | gacagctacgtgactacaga |
| gtgttaatttcttctcgtcc | tagccgaaaataccgactac | cgtcgaaaacgttcttgagc |
| caaaaagccggcgactagtc | attttcgtccgctatagaag | ctcccacataacggatgtaa |
| tgtcctttacttgattttct | gtgtttttaaactccacggc | caaactgcccttatagctga |
| cttgacctacttatggtgta | tcaatatggctcacgaatcc | gctaagcgtgctactcacat |
| ttttagttttgtggactcga | acgatctgtctctttatcag | ttgccacaacgtagctatgt |
| gtaaggaaaacagccgcctc | gccttgccaattatgtgcaa | cgaccgtaacaacgactaag |
| agcccatgaattcgatacta | atctttaggtaggctaacgg | ttctttcgacattatttcgg |
| ctcggaagacaaggaagcgt | gacttctactctctgacttc | gcgacgacttttgctaagta |
| ttgtctgtaccttttcacat | cgagtacttcctactttttc | aagtatcgttcctcctattt |

**Table S3. Sequences of fluorescently labelled probes (3′ to 5′ orientation) targeting *trpE* mRNA in (A) *E. coli* and (B) *B. subtilis*.** Probes were designed using the Stellaris Probe Designer (LGC Biosearch Technologies) with an oligonucleotide length of 20 nucleotides, a minimum spacing of 2 nucleotides, and a masking level of 1–2.
